# Clustering-based optimization method of reference set selection for improved CNV callers performance

**DOI:** 10.1101/478313

**Authors:** Wiktor Kuśmirek, Agnieszka Szmurło, Marek Wiewiórka, Robert Nowak, Tomasz Gambin

## Abstract

**Background:** There are over 25 tools dedicated for the detection of Copy Number Variants (CNVs) using Whole Exome Sequencing (WES) data based on read depth analysis.

The tools reported consist of several steps, including: (i) calculation of read depth for each sequencing target, (ii) normalization, (iii) segmentation and (iv) actual CNV calling. The essential aspect of the entire process is the normalization stage, in which systematic errors and biases are removed and the reference sample set is used to increase the signal-to-noise ratio.

Although some CNV calling tools use dedicated algorithms to obtain the optimal reference sample set, most of the advanced CNV callers do not include this feature.

To our knowledge, this work is the first attempt to assess the impact of reference sample set selection on CNV detection performance.

**Methods:** We used WES data from the 1000 Genomes project to evaluate the impact of various methods of reference sample set selection on CNV calling performance of three chosen state-of-the-art tools: CODEX, CNVkit and exomeCopy. Two naive solutions (all samples as reference set and random selection) as well as two clustering methods (k-means and k nearest neighbours with a variable number of clusters or group sizes) have been evaluated to discover the best performing sample selection method.

**Results and Conclusions:** The performed experiments have shown that the appropriate selection of the reference sample set may greatly improve the CNV detection rate. In particular, we found that smart reduction of reference sample size may significantly increase the algorithms’ precision while having negligible negative effect on sensitivity. We observed that a complete CNV calling process with the k-means algorithm as the selection method has significantly better time complexity than kNN-based solution.

## Background

Accurate detection of clinically relevant CNVs is essential in the diagnosis of genetic diseases since Copy Number Variants (CNVs) are responsible for a large fraction of Mendelian conditions [1, 2]. While the bioinformatics pipelines specializing in the detection of Single-Nucleotide Variants (SNVs) and short indels using WES data are mature and provide satisfactory performance https://precision.fda.gov/challenges/consistency, the identification of larger deletions and duplications still remains a challenge. Although a plethora of tools have been developed to call CNVs form WES data, most of these algorithms are characterized by limited resolution, insufficient performance, and unsatisfactory classification metrics [3, 4]. Although fine-tuning of CNV calling parameters may substantially improve the overall algorithm performance [5], there are still no well-established guidelines that would help optimize the detection rate of CNV-calling pipeline.

CNVs from WES can be detected based on the analysis of read-depth (RD) data. Typically, CNV calling algorithms can be broken down into four main stages. First, in the depth of coverage calculation step, the number of mapped reads in each exon is counted, followed by the quality control stage in which poorly covered exons and samples are removed. Next, the normalization process is applied to determine the depth of coverage under the assumption that CNVs do not occur. To minimize the effect of technological biases, CNV calling algorithms are required to take into account the depth of coverage in other samples (reference sample set) and the influence of known sources of noise, including but not limited to reads mappability and GC content in target regions. Then, the raw and normalized depth of coverage are compared - mainly the log ratio of raw depths of coverage versus normalized one is calculated. Finally, segmentation and actual CNV calling are applied, which produces a set of putative deletions and duplications.

The appropriate selection of the reference sample set has a substantial influence on the background modeling, and as a consequence, on the performance of a CNV caller. Unfortunately, most of the tools do not provide procedures for choosing the optimal reference set from among available samples except for CANOES [6], ExomeDepth [7] and CLAMMS [8]. The algorithm of selection in ExomeDepth and CANOES is based on counting correlation between the investigated sample and the rest of the samples aiming to find most similar elements and add them to the reference set. Then, k nearest neighbors (kNN) meaning the *k* most correlated samples are taken as a reference set for a specified element. In CANOES, the maximum number of samples in reference set is fixed and can be modified by the user (default 30), whereas in ExomeDepth this number is determined by the algorithm which maximizes the posterior probability in favour of a single-exon heterozygous deletion call by ExomeDepth’s model. In CLAMMS, the number of selected samples in the reference set is also set by the user. CLAMMS uses kNN algorithm as well, but, in contrary to CANOES and ExomeDepth, the distance metric between samples is counted basing on Binary Alignment Map (BAM) features (extracted by Picard (http://broadinstitute.github.io/picard)).

In this work, we investigated four different approaches to the selection of reference sample set labelled “all samples”, “random”, “kNN” and “k-means”. They were applied to the subset of 1000 Genomes data to study the influence of a selection method on the performance of CNV callers. In our analysis we combined reference set selection methods with three chosen state-of-the-art algorithms specializing in the identification of CNVs from WES data which do not provide any automated mechanism for selecting reference samples (CODEX, exomeCopy, and CNVkit).

## Methods

### Benchmark dataset

To evaluate the influence of the reference sample set selection on the CNV calling performance of selected algorithms, we used 1000 Genomes project phase 3 WES data from 861 individual (444 females and 417 males), including 205 samples from Europe (106 samples from Tuoscany in Italia; 99 Utah Residents (CEPH) with Northern and Western European Ancestry), 276 samples from Africa (109 samples from Yoruba in Ibadan, Nigeria; 101 samples from Luhya in Webuye, Kenya; 66 Americans of African Ancestry in SW USA), 207 samples from East Asia (103 samples from Han Chinese in Beijing, China; 104 samples from Japanese in Tokyo, Japan), 106 samples from South Asia (Gujarati Indian from Houston, Texas), and 67 samples from America (Mexican Ancestry from Los Angeles USA).

The investigated samples were sequenced by the seven research centres [9]. The correlation analysis of coverage profiles among the samples confirmed the existence of several clusters that correspond to different capture designs used in the project (Fig. 1).

**Figure 1.**
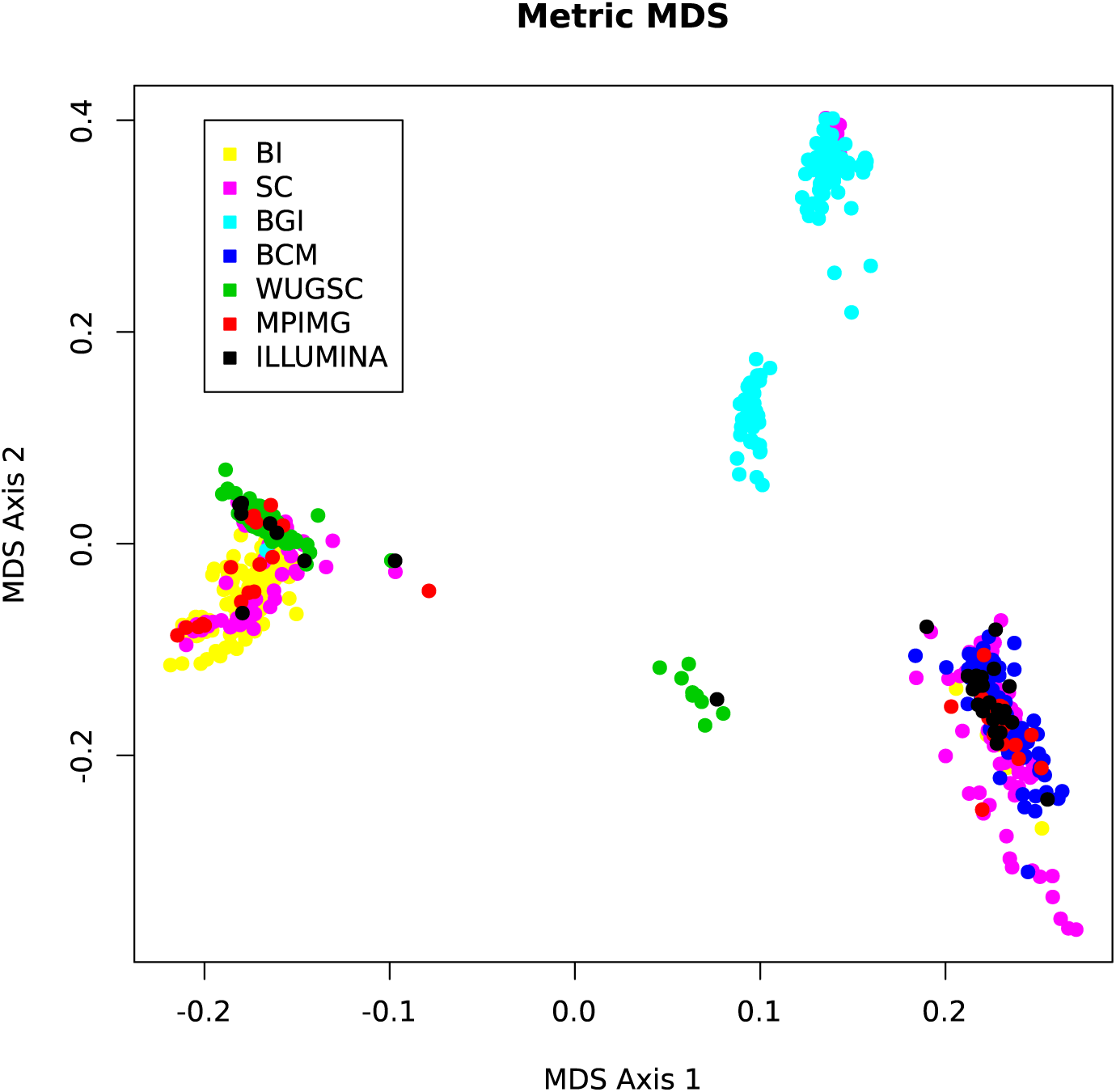
Correlation between samples of benchmark dataset. The figure presents the results of a multidimensional scaling of the covariance matrix of the read count data for the 861 investigated samples onto a two-dimensional plane. The colors depict samples from other sequencing centres (BCM - Baylor College of Medicine, BGI - Bejing Genomics Institute, BI - The Broad Institute, ILLUMINA - Illumina, MPIMG - The Max Planck Institute of Molecular Genetics, SC - The Welcome Trust Sanger Institute, WUGSC - Washington University Genome Science Center). It is worth noticing that samples are grouped into several clusters, mainly according to the research center where they were sequenced. However, samples sequenced in the same research center are also divided into subgroups, e.g. cyan dots, which depict the samples from Baylor College of Medicine. The figure was prepared by R’s cmdscale function.

The quality control (as implemented in CODEX) was performed for each sample. In this process all targets with median read depth across all samples below 20 or greater than 4000, targets shorter than 20 bp or longer than 2000 bp, with mappability factor below 0.9 and GC content below 20% or greater than 80% were removed. In our analysis we considered the first chromosome only in order to reduce the computation time.

### Study design

The presented approach follows fork-join [10] processing model with each sample being processed separately (possibly in parallel) and combining many outputs into the final CNVs collection. Operations performed on a single element include selecting the reference set, followed by normalization and CNV caller invocation, producing a list of detected variants for a considered sample. The union of all partial results creates the set of detected CNVs for the whole input sample set (Fig. 2).

**Figure 2.**
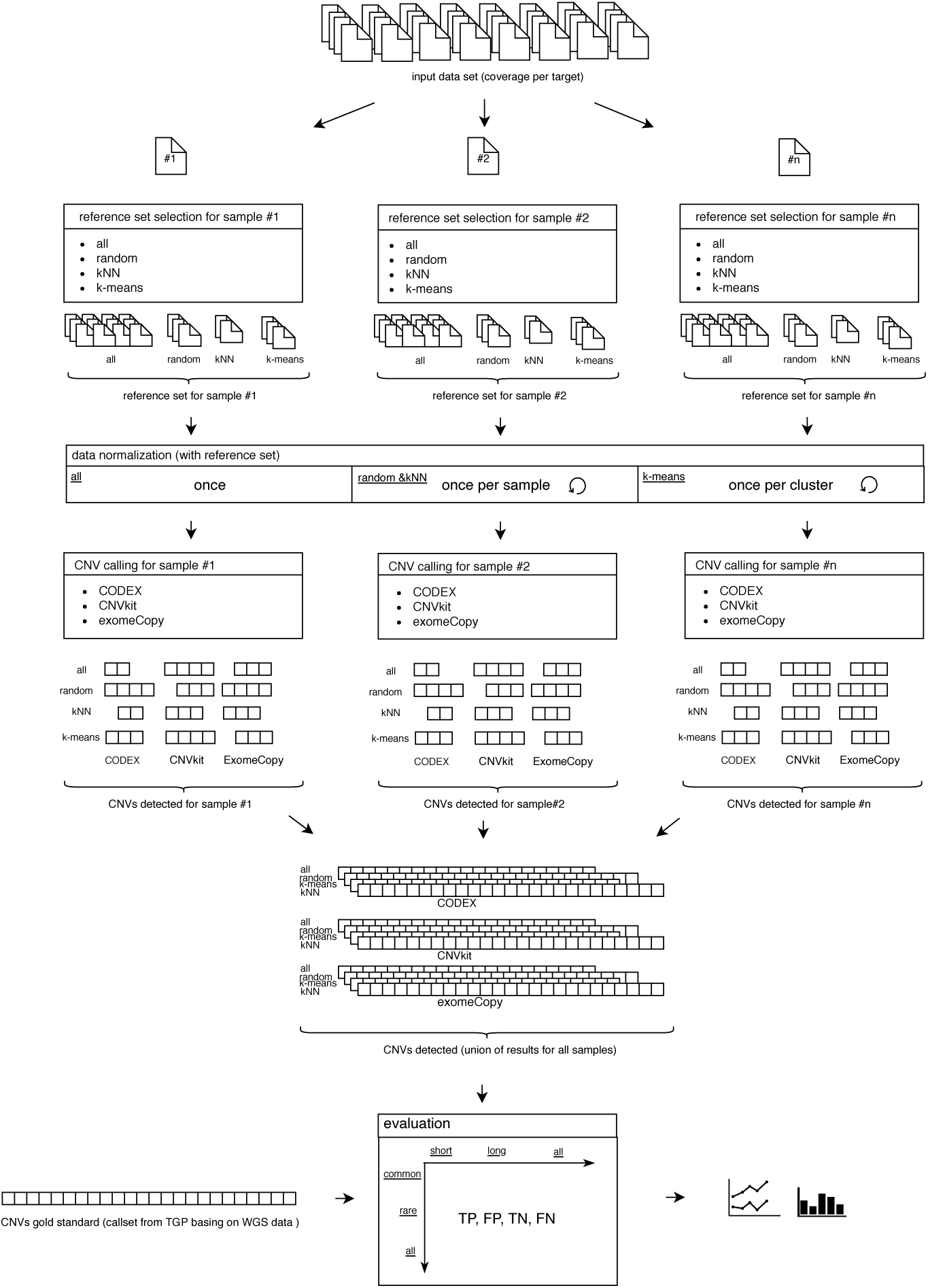
Workflow of the research method presented in the study. The input dataset is a set of numbers which depict the depth of coverage in samples on specified exons. Each sample from this dataset is processed by a reference sample set selector module, which is responsible for designating a set of samples that will be the reference collection. As a consequence, every element from the input dataset has its own, independent reference set. The normalization step uses the determined reference sets to perform normalization. This step is performed only once for “all” method, once per sample in “kNN” and “random” approach and once per cluster in “k-means” strategy. Then, to each generated sample, we apply CNV calling performed by three callers: exomeCopy, CODEX and CNVkit. The input for the CNV detecting tool is a set of samples consisting of the investigated sample and its reference panel. After calling CNVs, the events in the investigated sample are filtered and, this set of events is added to the final set of CNVs. Having processed all samples from the input dataset, the union of all partial per-sample results stored as the output call set for each approach combining selection method and variant caller. The evaluation of the results is performed against CNVs call set gold standard, delivered by 1000 Genomes Project. Variations are additionally categorized into common and rare as well as short and long categories, which allows us to precisely calculate the True Positives, True Negatives, False Positives and False Negatives metrics in those groups.

### Reference set selection

To asses the impact of reference set selection on the performance of a CNV caller, we have chosen and examined four approaches. The first method (”all samples”), considered as a baseline solution, encompasses all samples from the input dataset, with no selection performed.

The second one (”random”) is a naive approach to selection, which is an arbitrary subset choice, implemented as a draw with no repetitions. We iterated this experiment 10 times with various sizes of reference collection (from 50 to 500). The third method (”kNN”) is the approach used in previous works (i.e., Canoes, ExomeDepth, CLAMMS), where only *k* most similar samples are included in the reference set (k nearest neighbors algorithm [11]). The distance metric between the elements is based on Pearson correlation between read depth of samples. We repeated the experiment 10 times with varying sizes of the desired set, 50 ≤ *k* ≤ 500. Finally, we create a reference set by a new approach (”k-means”) using the k-means [12] clustering algorithm. The whole sample set is divided into *k* groups, basing again on the correlation between read depth of elements as proximity measure. We repeated the experiment 10 times with varying *k* value (from 1 to 10).

### Normalization and CNV calling

In our experiments, normalization of read depth data and CNV calling was performed by three state-of-the-art tools (CODEX v.1.8.0, exomeCopy v.1.22.0 and CNVkit v.0.9.3) that do not perform the reference set selection on their own.

CODEX algorithm is based on a multi-sample normalization model, which is fitted to remove various biases including noise introduced by different GC content in the analyzed targets. CNVs are called by the Poisson likelihood-based segmentation algorithm. ExomeCopy [13], on the other hand, implements a hidden Markov model which uses positional covariates, including background read depth and GC content, to simultaneously normalize and segment the samples into the regions of constant copy count. CNVkit [14] uses both the targeted reads and the non-specifically captured off-target reads to distribute the copy number evenly across the genome.

All of the tools were called with their default parameters, except for the one related to lowering the maximum numbers of the latent factors in CODEX (from 9 to 3) to reduce the calculation time and the changing default segmentation method in CNVkit application from default circular binary segmentation to HaarSeg, a wavelet-based method [15]. For the “all” and “random” approach the normalization step is called once, followed by CNV calling for each sample. “kNN” method results in a different reference set for each sample; therefore, it requires both a normalization and a calling stage performed for each input element. With the “k-means” method it is sufficient to normalize samples only once per group, followed by the CNV calling step.

### Performance evaluation

We have evaluated the quality of each pair of (i) reference set selection algorithm and (ii) CNV calling tool, comparing the output CNV call set of the solution and the CNV call set golden record provided by 1000 Genomes Consortium [9] generated based on the Whole Genome Sequencing (WGS) data. To asses accurately the influence of the reference set on the final output, the results were evaluated separately for different subset of CNVs. The variants were categorized into rare (frequency ≤ 5%), common (> 5%) CNVs, as well as short (encompassing 1 or 2 exons) and long (encompassing more than 3 exons) CNVs. As part of the evaluation stage we calculated the Dunn Index [16] and Silhouette width [17] to discover the quality of k-means clustering for varying value of k. The above mentioned measures are based solely on grouped data, presenting to what extent the clusters formed are compact and well separated.

## Results

### All samples as reference set

The “all samples” strategy is treated as a baseline for further evaluation and is equivalent to the default mode of CNV callers invocation without any selection. Note, that the same reference set is generated by the kNN algorithm in case of k equal to the number of all samples and by k-means with single cluster (*k* = 1). Overall, we found CNVkit to have the highest precision but the lowest sensitivity among the examined callers. The F1 measure was the highest for CODEX (0.34), followed by CNVkit (0.32) and exomeCopy (0.05).

### Random reference set

The performed experiments revealed that random selection of the reference sample set does not significantly affect the number of true-positive and false-positive calls. Interestingly, the performance statistics do not change significantly as the number of random samples in the reference panel increases for all of the CNV callers (Fig. 3A, B).

**Figure 3.**
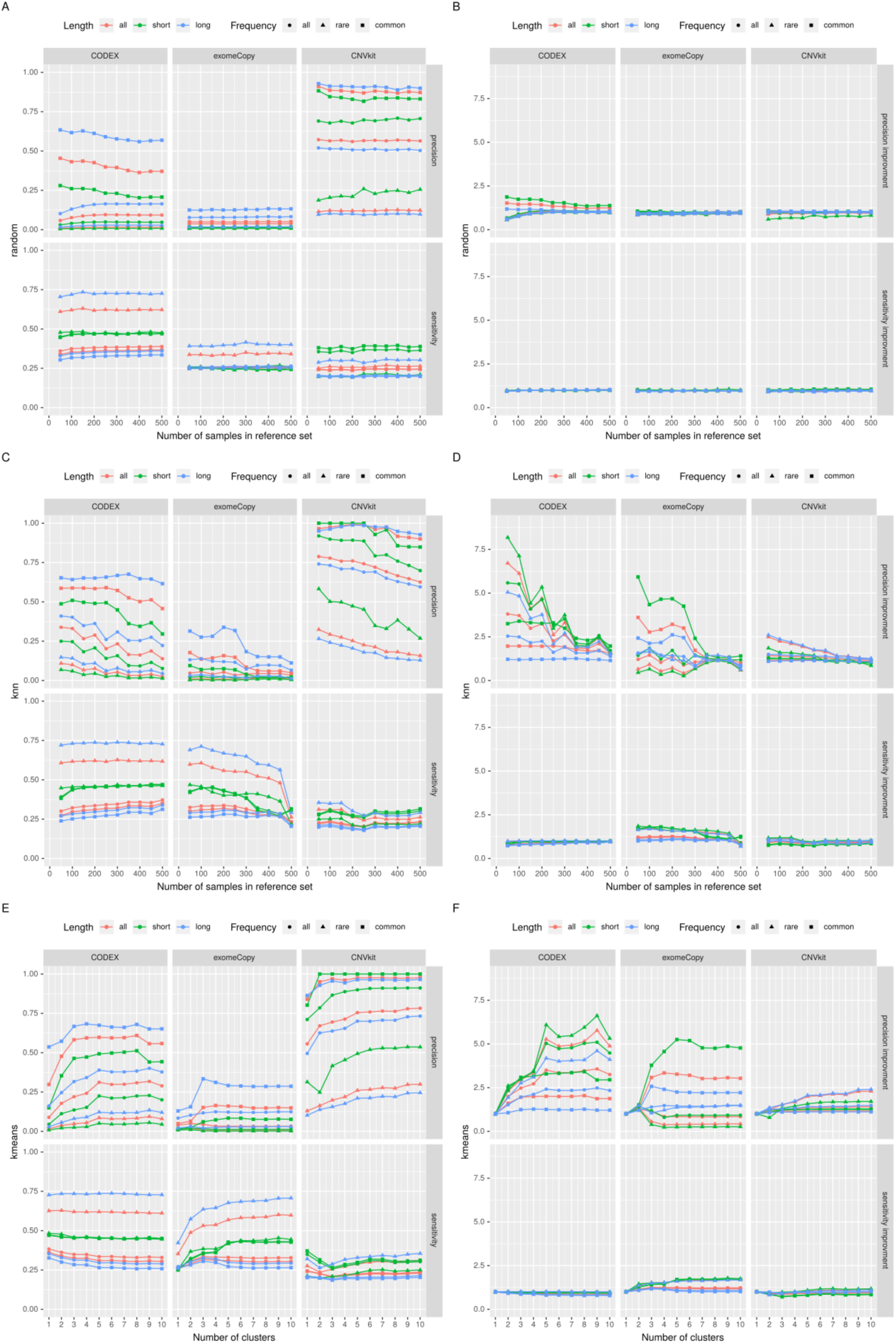
Results of four selection method used in CODEX, CNVkit and exomeCopy. Panels A, C, E present absolute changes in the precision and sensitivity of the investigated CNV callers for different methods of the reference set selection; relative performance in relation to baseline is presented in panels B, D, and F. The results for the “all” method (baseline) are presented in the “kmeans” diagram, where k is equal to 1 (single group).

### k Nearest Neighbors

We found that sensitivities of CODEX and CNVkit callers are independent of the *k* value in the k Nearest Neighbors algorithm, whereas the sensitivity of exomeCopy is inversely proportional to *k*, especially, for rare events (Fig. 3C, D). On the other hand, the precision of exomeCopy is rather stable, whereas in case of CODEX and CNVkit the precision increases when *k* is growing.

### k-means

The results show that the precision for CODEX and CNVkit is significantly greater, when *k* ≥ 5 in comparison to a single group (see Fig. 3E, F). We observed that for both tools the saturation point occurs at *k* = 4 or *k* = 5. This point represents the optimal, minimal number of groups and it is in line with both: (i) the number of groups that emerge on the Fig. 1 and (ii) the optimal values of Dunn Index and Silhouette width (Fig. 4). Sensitivity for CODEX and CNVkit remains fixed for different numbers of clusters in the k-means algorithm for *k* exceeding 5. exomeCopy demonstrates opposite characteristics - constant precision, and significantly improved sensitivity for a number of groups greater or equal to 5 in comparison to a single group, especially, for short calls.

**Figure 4.**
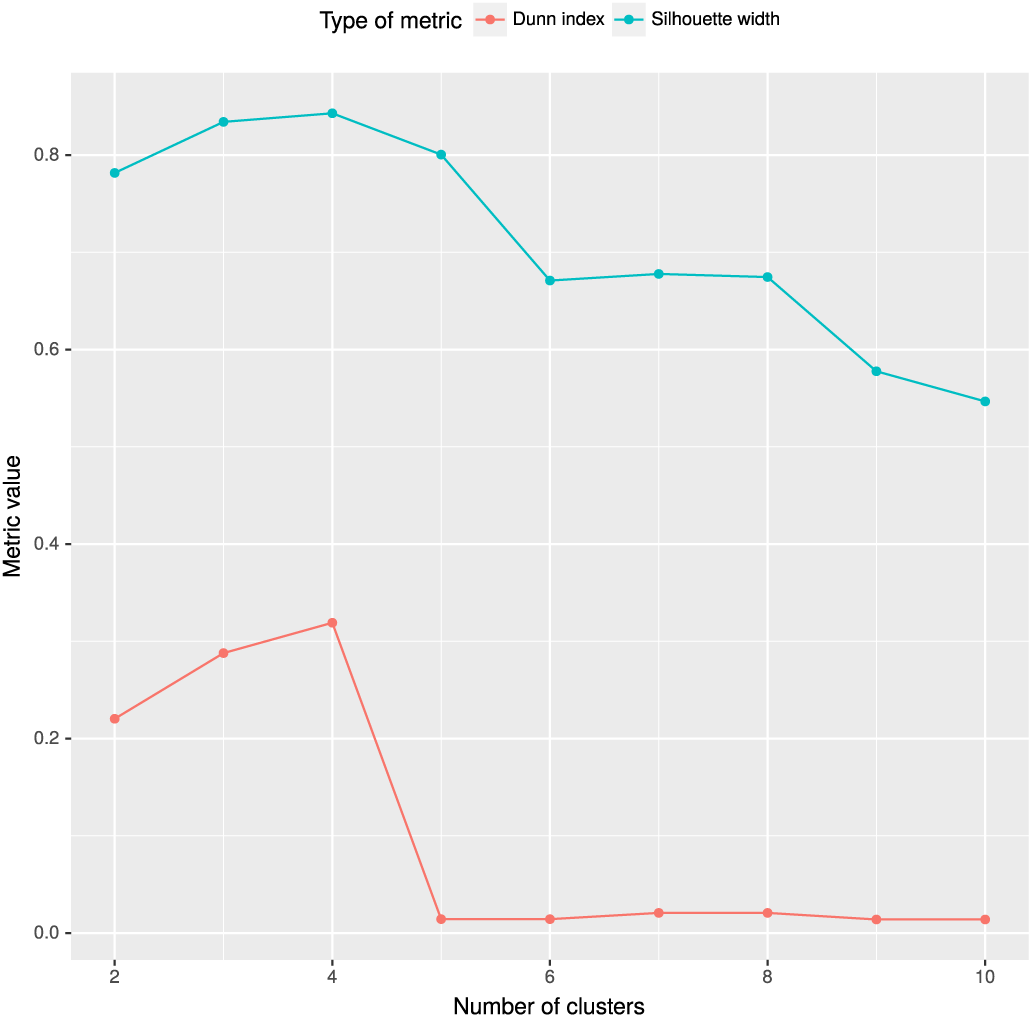
Dunn Index and Silhouette width for assessing the number of groups in k-means algorithm. The figure presents the evaluation of the number of groups by means of two metrics, which combine the measures of compactness and separation of the clusters. Briefly, the higher the value of both indexes, the better the division into clusters. The figure shows that for the data set examined in the presented work, the most optimal number of groups is 4, which agrees with the figure of sensitivity and precision - for example, precision for CODEX tool.

### Comparison of reference set selection methods

CODEX achieved the highest F1 score in our benchmark; hence we used evaluation results from this algorithm to compare different methods of reference set selection (Fig. 5). The results for “kNN” and “k-means” approaches differ depending on *k* value. For comparison we have chosen the best performing *k* values for both approaches. In “k-means” case, *k* is equal to 4, according to internal quality measures (Fig. 4). In “kNN”, *k* equals the value of the quotient of a total number of samples and the best performing number of clusters resulting in *k* ≈ 200. The analysis confirmed that CNV calling performance was much higher when “kNN” or “k-means” approaches were used instead of “all samples” or “random” methods. Interestingly, we observe essentially no difference in CNV calling accuracy between “kNN” and “k-means”. This is a particularly important conclusion, since the latter method is characterized by better time complexity than the former one (Fig. 6).

**Figure 5.**
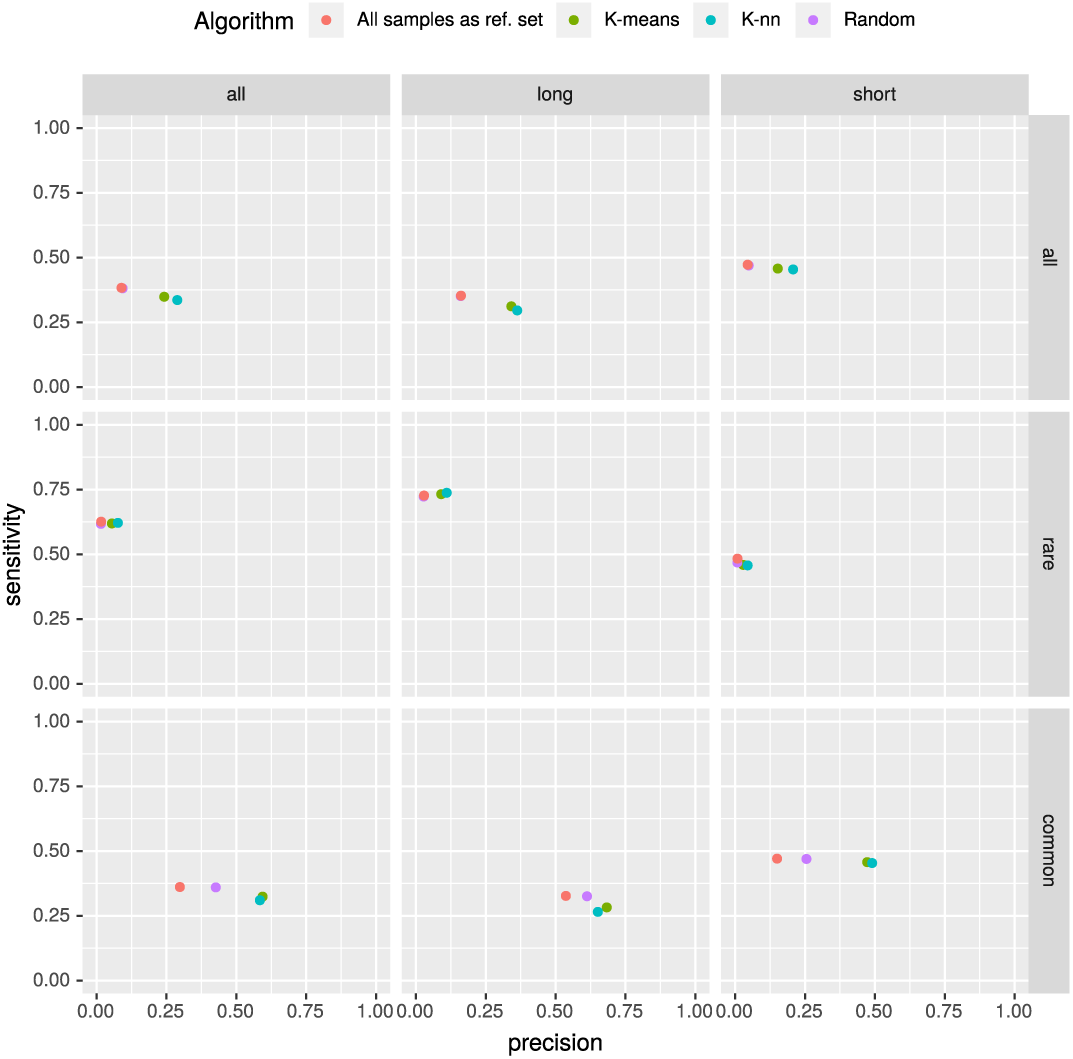
Comparison of CNVs detected by CODEX with four different selection methods. The k value for “k-means” method was equal to 4, for “kNN” algorithm - 200; the baseline, “all samples” method, is distinguished as “k-means” approach for *k* = 1. The figure shows, that using “kNN” and “k-means” methods results in better precision in comparison to the baseline. What is more, sensitivity for all of the investigated methods remains fairly stable. It is worth noting that the dots for the “kNN” and “kmeans” methods are very close to each other - both of mentioned methods lead to very similar results.

**Figure 6.**
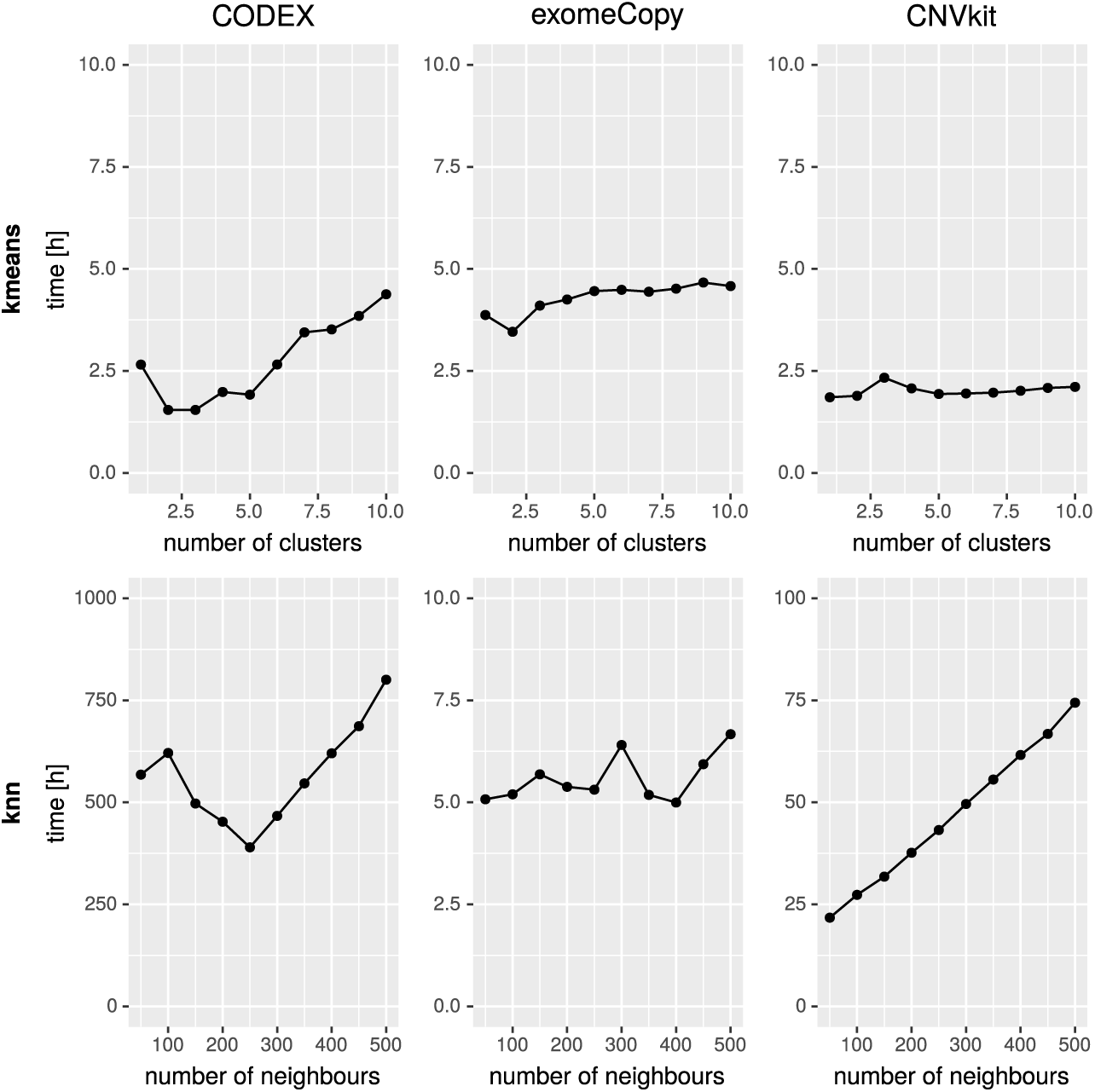
Comparison of CNV calling detection time in large cohort of samples using “k-nn” and “k-means” based approach to reference selection. The time measured includes reference set selection, normalization and CNV calling step. Note that CNV calling is performed for all samples in the cohort. The substantial difference between total execution times is a consequence of normalization process as required by kNN method (once per sample) and k-means approach (once per cluster). Since the number of samples is significantly greater than the number of groups, the total normalization time is significantly greater for “kNN” than for “k-means”.

## Discussion

### Performance of CNV callers without reference sample set selection

The sensitivity and precision of CNV callers vary owing to different approaches being implemented in those tools (see Fig. 3). As expected, CNV detecting solutions are more sensitive in discovering rare events, whereas large and common CNVs are especially difficult to be detected since it is hard to distinguish them from a common technical artifacts and biological biases. At the same time, our results indicated that the most challenging issue is related to a very high number of false positive calls observed in the class of rare variants. In this context, one of the most important finding in our study is that the precision in calling rare CNVs (i.e., reduction of the number of false positives) could be significantly improved by the application of “k-nn” or “k-means” based methods of reference set selection.

### Random selection of reference set does not change the CNV calling performance

The naive approach to selecting the reference set as a random subset mostly does not change the CNV callers’ characteristics. Since this approach does not positively impact the performance, it is not recommended.

### Appropriate selection of reference samples improves CNV detection

In this paper we have shown that the correct reference set selection improves the results of all tested CNV callers. The highest improvement was achieved for the class of short CNVs, which are usually the most challenging to be identified and often missed by orthogonal biological assays, including Comparative Genomic Hybridization (CGH) arrays. In case of CODEX, the precision of short, rare CNVs detection increased more than seven times when using “k-means” or “kNN” in comparison to “all” or “random” strategies. Moreover, sensitivity of exomeCopy in detecting short, rare CNVs was more than two times greater when clustering-based or “kNN” strategies are used. These findings are particularly important since the more accurate detection of rare and short CNVs may substantially improve the molecular diagnostic solution rate in clinics. The main aim of selecting the correct reference set is determining the most similar samples. In order to identify the best performing number of clusters in k-means algorithms, we have used Silhouette and Dunn measures

### k-means vs. kNN based approach

The experiments proved that performance metrics for reference sets chosen by kNN and k-means methods are similar. Although the existing tools (Canoes, ExomeDepth, CLAMMS) use kNN-based methods as the reference set selection algorithm in this study, we have shown that k-means has much less time complexity. As Figure 6 clearly states, the “kNN” approach is significantly (approximately 200x) slower. The reason for this is the need to invoke a long-lasting normalization process as many times as the number of input elements is, while “k-means” requires normalization only once per cluster.

The reference set selection with the k-means algorithm could be performed on any data set by splitting samples on a given number of groups with simultaneous monitoring of Dunn Index and Silhouette width values.

### Further research

The solution that would automatically determine, based on the input data set, the best parameters for CNV calling would facilitate the entire process. To achieve this, we are designing a module for the injection of artificially generated CNVs into the user data and the comparison of the detected CNVs with the events injected. This technique will enable the choice of an appropriate method for the reference sample set selection for any WES data set as well as automatic selection of parameters of a given method for the selection of reference sample set including the number of clusters in the k-means algorithm. This process could be included in complex tools, e.g. applications for optimization CNV callers like Ximmer [5].

## Conclusions

We have shown that proper reference sample set selection leads to improved sensitivity and precision for all considered CNV callers. Our results revealed that k-means and kNN based approaches guarantee a very similar performance of CNV calling while the former one is significantly faster and requires less computational resources. Finally, we have shown that the optimal number of groups for k-means algorithm (corresponding to the highest accuracy of CNV calling) can be estimated using internal clustering metrics (such as Dunn Index and Silhouette width). To summarize, these conclusions can be used as a guideline that helps in appropriate implementation and fine-tuning of CNV calling pipelines from WES data in the clinical and research environment.

## Abbreviations

Not applicable.

## Declarations

## Availability and requirements

### Authors’ contributions

TG identified the problem, WK and TG designed the approach. WK implemented the software. WK and TG worked on testing and validation. WK, TG, AS, RN, MW wrote the manuscript. All authors read and approved the final manuscript.

### Ethics approval and consent to participate

Not applicable.

### Consent for publication

Not applicable

### Additional files

Not applicable.

### Author details

^1^ Institute of Computer Science, Warsaw University of Technology, ul. Nowowiejska 15/19, 00-665 Warsaw, Poland.

^2^.

## Copyright

© The Author(s) 2018

## References

1. Zhang, F., Gu, W., E Hurles, M., R Lupski, J.: Copy Number Variation in Human Health, Disease, and Evolution 10, 451–81 (2009)

2. Stankiewicz, P., R Lupski, J.: Structural Variation in the Human Genome and its Role in Disease 61, 437–55 (2010)

3. Yao, R., Zhang, C., Yu, T., Li, N., Hu, X., Wang, X., Wang, J., Shen, Y.: Evaluation of three read-depth based cnv detection tools using whole-exome sequencing data. Molecular Cytogenetics 10(1), 30 (2017). doi:10.1186/s13039-017-0333-5

4. Tan, R., Wang, Y., Kleinstein, S.E., Liu, Y., Zhu, X., Guo, H., Jiang, Q., Allen, A.S., Zhu, M.: An evaluation of copy number variation detection tools from whole-exome sequencing data. Human Mutation 35(7), 899–907. doi:10.1002/humu.22537. https://onlinelibrary.wiley.com/doi/pdf/10.1002/humu.22537

5. Sadedin, S., Ellis, J., Masters, S., Oshlack, A.: Ximmer: A system for improving accuracy and consistency of cnv calling from exome data. bioRxiv (2018). doi:10.1101/260927. https://www.biorxiv.org/content/early/2018/02/06/260927.full.pdf

6. Backenroth, D., Homsy, J., Murillo, L.R., Glessner, J., Lin, E., Brueckner, M., Lifton, R., Goldmuntz, E., Chung, W.K., Shen, Y.: Canoes: detecting rare copy number variants from whole exome sequencing data. Nucleic Acids Research 42(12), 97 (2014)

7. Plagnol, V., Curtis, J., Epstein, M., Y Mok, K., Stebbings, E., Grigoriadou, S., Wood, N., Hambleton, S., Burns, S., J Thrasher, A., Kumararatne, D., Doffinger, R., Nejentsev, S.: A robust model for read count data in exome sequencing experiments and implications for copy number variant calling 28, 2747–2754 (2012)

8. S Packer, J., K Maxwell, E., O’Dushlaine, C., E Lopez, A., E Dewey, F., Chernomorsky, R., Baras, A., D Overton, J., Habegger, L., G Reid, J.: Clamms: A scalable algorithm for calling common and rare copy number variants from exome sequencing data 32 (2015)

9. The 1000 Genomes Project Consortium: A global reference for human genetic variation. Nature 526, 68–74 (2015)

10. Conway, M.E.: A multiprocessor system design. In: Proceedings of the November 12-14, 1963, Fall Joint Computer Conference. AFIPS’63 (Fall), pp. 139–146. ACM, New York, NY, USA (1963). doi:10.1145/1463822.1463838. http://doi.acm.org/10.1145/1463822.1463838

11. Zhang, Z.: Introduction to machine learning: K-nearest neighbors. Annals of Translational Medicine 4, 218–218 (2016)

12. McQueen, J.: Some methods for classification and analysis of multivariate observations 4, 257–272 (1967)

13. I. Love, M., MyÅ¡iÄkovÃ¡, A., Sun, R., Kalscheuer, V., Vingron, M., A. Haas, S.: Modeling read counts for cnv detection in exome sequencing data 10, 52–52 (2012)

14. Talevich, E., Shain, A., Botton, T., C Bastian, B.: Cnvkit: Genome-wide copy number detection and visualization from targeted dna sequencing 12, 1004873 (2016)

15. Ben-Yaacov, E., C Eldar, Y.: A fast and flexible method for the segmentation of acgh data 24, 139–45 (2008)

16. C. Dunn, J.: A fuzzy relative of the isodata process and its use in detecting compact well-separated clusters. Cybernetics and Systems 3, 32–57 (1973). doi:10.1080/01969727308546046

17. Rousseeuw, P.J.: Silhouettes: A graphical aid to the interpretation and validation of cluster analysis. Journal of Computational and Applied Mathematics 20, 53–65 (1987). doi:10.1016/0377-0427 (87)90125-7

